# Antagonistic effects of chemical mixtures on the oxidative stress response are silenced by heat stress and reversed under dietary restriction

**DOI:** 10.1101/2021.03.17.435857

**Authors:** Karthik Suresh Arulalan, Javier Huayta, Jonathan W. Stallrich, Adriana San-Miguel

## Abstract

Chemical agents released into the environment can induce oxidative stress in organisms, which is detrimental for health. Although environmental exposures typically incorporate multiple chemicals, organismal studies on oxidative stress derived from chemical agents commonly study exposures to individual compounds. In this work, we explore how chemical mixtures drive the oxidative stress response under various conditions in the nematode *C. elegans*, by quantitatively assessing levels of *gst-4* expression. Our results indicate that naphthoquinone mixtures drive responses differently than individual components, and that altering environmental conditions, such as increased heat and reduced food availability, result in dramatically different oxidative stress responses mounted by *C. elegans*. When exposed to heat, the oxidative stress response is diminished. Notably, when exposed to limited food, the oxidative stress response specific to juglone is significantly heightened, while identified antagonistic interactions between some naphthoquinone components in mixtures are abolished. This implies that organismal responses to xenobiotics is confounded by environment and stressor interactions. Given the high number of variables under study, and their potential combinations, a simplex centroid design was used to capture such non-trivial response over the design space. This makes the case for the adoption of Design of Experiments approaches as they can greatly expand the experimental space probed in noisy biological readouts, and in combinatorial experiments. Our results also reveal gaps in our current knowledge of the organismal oxidative stress response, which can be addressed by employing sophisticated design of experiments approaches to identify significant interactions.

## Introduction

Oxidative stress, which has deleterious effects on health^1–3^, can be induced by ROS (Reactive Oxygen Species) generated from oxidant chemicals. The effects of oxidant exposures on biological systems have been an important area of study, although these effects have been mostly analyzed by employing chemical exposures of individual components.^4^ However, realistic environmental exposures are mixtures of multiple components.^5–7^ In addition, environmental factors, such as diet and temperature, can modulate the mechanisms by which chemicals induce toxicity and activate defense responses in living organisms.^7,8^ It is still unclear how chemical mixtures drive oxidative stress and how environmental conditions modify such responses. In this work, we analyze how mixtures of oxidant species, in particular naphthoquinones, differentially drive the oxidative stress response in the model organism *C. elegans*.

Naphthoquinones are strong oxidants. They are used as precursors of toxic industrial chemicals.^9^ They are also a product of fossil fuel combustion and atmospheric photochemical conversions, and are thus found in ambient particulate matter.^10^ Environmental exposures to naphthalene (a precursor for naphthoquinones) are significant as it is found in the atmosphere and in cigarette smoke.^11–13^ Naphthalene toxicity is thought to occur through the action of napthoquinones.^14,15^ Naphthoquinones have been hypothesized to induce oxidative stress through two distinct mechanisms^16,17^: depletion of glutathione (an antioxidant that neutralizes ROS and counteracts xenobiotics by conjugation) through Michael reaction, or production of ROS through redox cycling.^16^ It is unclear if differences in the cytotoxic mechanisms amongst naphthoquinones could be reflected as differences in organismal responses to naphthoquinone mixtures. Naphtoquinone exposures at low concentrations can also induce beneficial effects. Simultaneous exposure to naphthoquinone derivatives such as juglone and plumbagin drive SKN-1/NRF-2 transcription factor-mediated hormesis in *C. elegans* at low concentrations, but turn toxic at higher concentrations.^18^ Naphthoquinones have also been shown to have anti-inflammatory effects in other model organisms.^19^ For example, allergen induced rats treated with juglone had a reduction in pulmonary eosinophils and bronchoalveolar lavage fluid.^20^ Given their prevalence as derivatives of naphthalene, and the differential responses to naphthoquinone exposures, it is critical to study the effects of naphthoquinone mixtures on organismal health.

The model system *C. elegans* facilitates studies on toxicity in a live organism with a well characterized nervous system, cell lineage, and physiology. *C. elegans* has proven to be a powerful model due to its small size, easy maintenance, and mapped genome and neuronal wiring. It enables *in vivo* studies using fluorescent markers due to their transparent bodies and ease of genetic manipulation.^21^ *C. elegans* has also been useful to study chemical mixture toxicity through growth and fertility assays.^22–25^ *C. elegans* and humans share concordant pathways, such as the insulin/IGF-1 signaling pathway (IIS), which regulates lifespan and healthspan extension driven by dietary restriction.^26^ Two-thirds of human proteins have homologs in *C. elegans*.^26^ *C. elegans* has been used to study mechanisms of toxicity identification, such as those produced by phorbol esters.^26^ Finally, toxicological studies have shown up to 69% concordance between *C. elegans* and mammalian toxicological data.^27^ Another study identified concordance between *C. elegans* data and that from rabbits and rats to be in the 45-53% range, just slightly lower than the concordance between rabbit and rat data (58%).^28^

Defense mechanisms to metabolize and eliminate xenobiotics are evolutionarily conserved from single cell organisms to humans.^29^ In mammalian cells, NRF-1, NRF-2, NRF-3 are a class of NF-E2-related factor 2 (NRF) transcription regulators employed for such defense mechanisms.^30^ In *C. elegans,* the oxidative stress response pathway is activated to counteract the toxicity caused by oxidative stressors.^31^The SKN-1 transcription factor, the functional ortholog of mammalian NRF-2, is major regulator of the oxidative stress response in *C. elegans*.^30,32,33^ It should be noted that SKN-1 diverges from NRF-2 in the way it binds to DNA. However, the similarities allow us to study SKN-1 in *C. elegans* as a model for mammalian NRF-2.^30^ SKN-1 has also been linked to other much broader homeostatic functions such as reducing stress, counteracting lipid accumulation, mitochondrial biogenesis and mitophagy, among others.^30^ SKN-1 activity in response to chemicals and heavy metals has been studied by examining expression of SKN-1-regulated genes using endogenously expressed fluorescent reporters. SKN-1 driven expression is mostly studied in the intestine, where digestion and detoxification occur.^4,34–36^ Multiple antioxidant response elements (AREs) containing genes are downstream targets of SKN-1, such as gst-4 and gcs-1, which encode for drug-metabolizing glutathione S-transferase (GST-4) and gamma-glutamyl cysteine synthetase (GCS-1), important in glutathione synthesis, respectively. For example, prior work by Crombie et al. focused on studying the effect of environmental factors on the oxidative stress response to juglone using a two level full factorial design, by monitoring *gst-4* expression.^37^ *gst-4* is commonly used as proxy for SKN-1 activity and thus activation of the oxidative stress response.^38–40^

In this study, we analyzed the effects of naphthoquinone mixtures on the *C. elegans* oxidative stress response under various environmental conditions, as determined by a *gst-4* translational fluorescent reporter. We follow a Design of Experiments (DoE) approach that enables systematic examination of the input factors to determine the individual and combinatorial influence on the measured response,^41,42^ while avoiding the unfeasible number of experiments required for full factorial designs with multi-level factors. This approach minimizes the number of experimental runs necessary to measure interactions between three naphthoquinones and two environmental conditions, while enabling comparisons from independent biological populations. We quantify response surfaces for these ternary mixtures under different conditions. We find that naphthoquinone mixtures drive antagonistic interactions, but these interactions are drastically modified by dietary restriction. On the other hand, heat stress abolishes oxidative stress response to both individual components and mixtures of naphthoquinones. This *gst-4* response is independent of the IIS factor DAF-16. The ROS levels and resistance to acute oxidative stress that result from exposure to naphthoquinone mixtures are also modulated by environmental factors, in the opposite direction as *gst-4* expression.

## Materials and methods

### Strain maintenance

*C. elegans* was maintained on standard Nematode Growth Medium (NGM) plates seeded with OP50 *E. coli* bacteria and maintained at 25 ºC. Worms at day 1 of adulthood were bleached and age synchronized using standard protocols,^43^ and grown to the young adult stage before application of oxidative stress. The strains used in the experiments were CL2166: *dvLs19* [pAF15 (gst-4P:GFP::NLS)], MAH97 *muIs109* [daf-16p::GFP::DAF-16 cDNA + odr-1p::RFP], and N2 (wild type) which were obtained from the Caenorhabditis Genetics Center. Animals were exposed to either control (S-Medium) or naphthoquinone mixtures in liquid culture in the presence of HB101 *E. coli* as a food source, based on established procedures.^44^ *E. coli* was grown in LB media in the presence of 4 mM streptomycin. Bacteria were washed thrice with SB media, pelletized, and resuspended in S-Medium at a concentration of 100 mg mL^−1^. Bacteria was then killed by heat treatment at 65 °C for a period of 45 mins,^37^ to avoid bacterial metabolism to confound results.

### Chemical preparation

Juglone, 1, 4-naphthoquinone, and plumbagin (**Figure 1A**) were sourced from Fisher Scientific and stored according to supplier guidelines. 100 mM stock solutions were prepared for 1, 4-naphthoquinone and plumbagin by dissolving the powdered compounds in DMSO and stored at −20 °C. Juglone stock was prepared fresh before the start of each experiment, given its low stability.^4^

**Figure 1:**
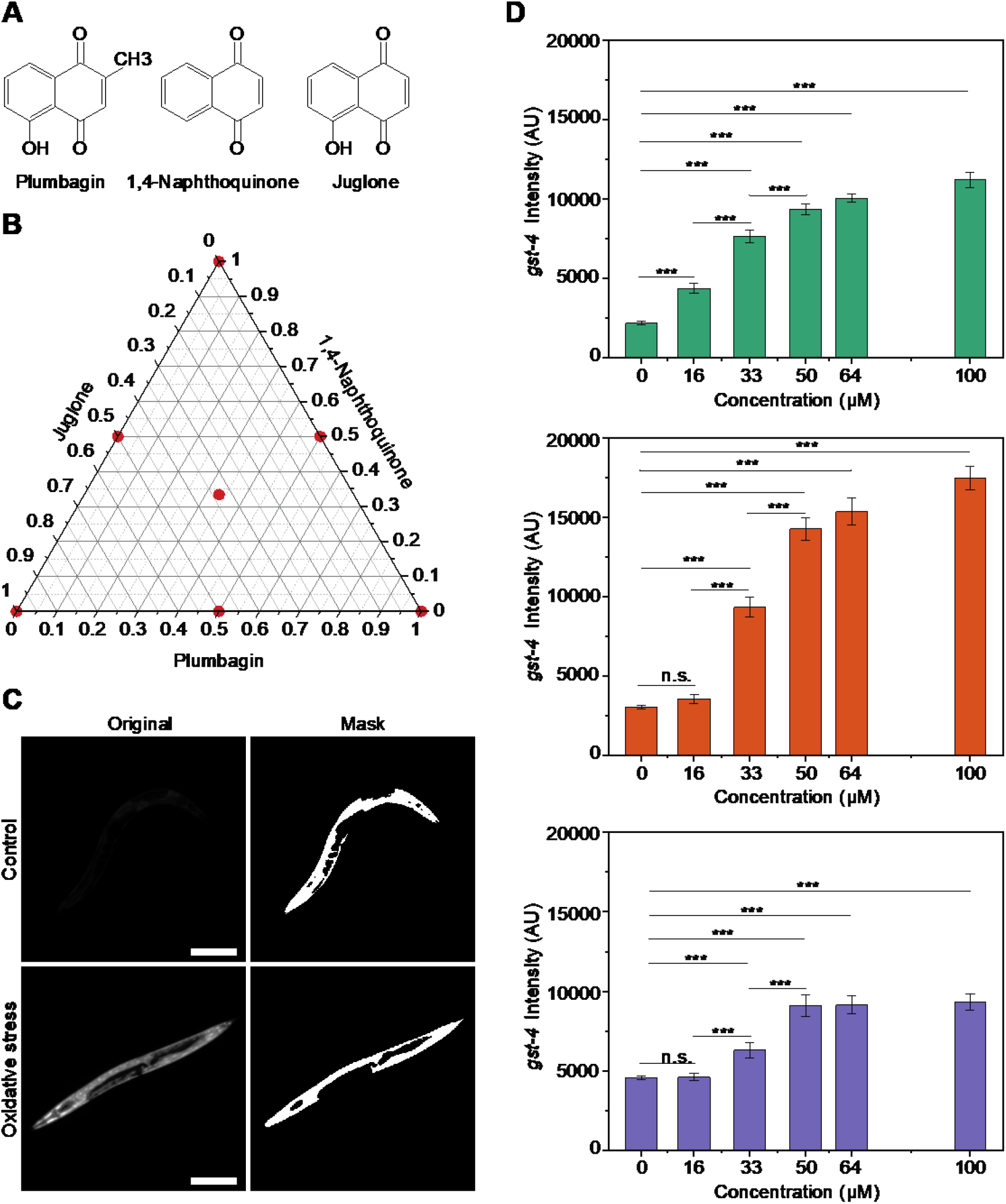
*gst-4* response is dose dependent. A) Compounds used for oxidative assays. B) Simplex centroid design. C) CL2166 worms under control and oxidative stress with corresponding masks for extracting *gst-4* expression levels. D) *gst-4* dose-dependent response to Plumbagin, 1, 4-Naphthoquinone, and Juglone (top to bottom). p > 0.05 (n.s.), p < 0.001 (***). Error bars are SEM. All p-values were calculated using Tukey HSD for all pairwise comparisons after one-way ANOVA (Unequal Variances) comparison in JMP 14.2. Scale bars are 200 μm.

### Experimental design

An experimental design was generated to study the effect of oxidant mixtures at *ad libitum* feeding (2 × 10^10^ cells mL^−1^) and no heat stress conditions, using a simplex centroid design (7 points in the experimental space, **Figure 1B**). The simplex centroid design was chosen as it entails a low number of experimental points, thus facilitating performing more replicates. This design entailed 40 runs split into 10 groups of 4 runs (where each design point had 5 – 7 replicates). To assign simplex centroid points to groups, we started from an I-optimal design generated in JMP Pro v 14.2 with 21 points in the experimental space, which already assigns each point to a group. These I-block points were then approximated to the closest point in the simplex centroid design, and their assignment to a group from the I-block design was maintained. An additional control run was tested along with each group to account for differences in populations due to any experimental variability (ambient temperature, humidity, etc.). A population of animals from a single NGM plate was thus divided into 5 runs (4 for design points, and one as a control).

To study the effects of process conditions on responses induced by chemical mixtures, a split plot mixture experiment was performed. The whole plot factors were the two environmental stressors and the split plot factors were the naphthoquinone mixtures. Two levels of heat exposures: 20 °C and sublethal 33 °C were considered as the first environmental stressor. Two levels of HB101 bacterial diet regime -*ad libitum* (2 × 10^10^ cells mL^−1^) (AL) and dietary restriction (2 × 10^9^ cells mL^−1^) (DR) was considered as the second environmental stressor.^37,45^ A total of 28 populations of animals were used for the split plot mixture experiments. Each population was split into five runs. One run, at process conditions 20 °C and *ad libitum* diet named “overall control”, was used to compare the 28 populations (7 points of simplex centroid design * 4 process conditions). Another run named “experimental control” was reserved for testing the effect of individual environmental stressor, i.e., a specific heat level and diet combination with no oxidant. Thus, each of the four process conditions were replicated 7 times across the 28 populations. The remaining three runs were exposed to the same combination of diet and heat exposure as the experimental control, but each was exposed to a different chemical mixture. These were assigned to groups similar to the previous experiment, using experimental points from a simplex centroid design. Hence, a total of 84 runs (7 points of the simplex centroid design * 3 replicates * 4 process conditions) and 56 control runs were used to study the effects of heat stress and dietary restriction on the naphthoquinone mixtures. The overall controls were plotted as Xbar and S control chart to ensure that populations were statistically comparable with each other (**Figure S1**).

### Application of oxidative stress and fluorescence imaging

Approximately 30-40 age-synchronized worms were loaded into a well of a 24-well plate containing the oxidant and HB101 bacteria in S-Medium. Control experiments were conducted with an empty vector of 1% v/v DMSO. Animals were exposed to the oxidants for a period of 8 hours at a temperature of 20 ± 1 °C. For Pgst-4::GFP imaging, worms were immobilized using a drop of 4 mM tetramisole on dried 2% agarose pads. Worms were imaged using a wide field inverted fluorescence microscope Leica DMi8 at 10x magnification after the exposure period. A Spectra X LED illumination system centered at 470 nm was used for excitation, and a Hamamatsu Orca Flash 4.0 16-bit digital CMOS camera was used for image acquisition. A dose response to individual compounds was first quantified by exposing animals in increasing concentrations from 0 μM to 100 μM for a period of 8 hours. The naphthoquinone mixtures were applied based on the designs described the previous section.

### Quantitative image processing

GFP driven by the *gst-4* promoter is observed throughout the animal, and its intensity was quantified to estimate oxidative stress response *in vivo*. Images were analyzed using a MATLAB script that generates a binary mask to identify a single worm per image. The MATLAB regionprops function was then used to quantify the mean intensity of the animal by overlapping the binary mask over the original image (**Figure 1C**).

### RNAi by feeding

To measure the influence of *daf-16* on the *gst-4* response to environmental conditions, age-synchronized worms were grown from the egg stage to day 1 of adulthood at 20 °C in NGM plates containing HB101 bacteria containing the dsRNA-producing vector from the Ahringer library (acquired from Source Biosciences).^46,47^ Animals were then bleached following standard protocols.^43^ The eggs obtained were deposited in new NGM plates containing HB101 bacteria carrying the *daf-16* RNAi vector. Animals were grown to young adulthood at 20 °C, and then exposed to oxidative stress as previously described.

### ROS detection

To measure ROS levels, 30 animals, exposed to oxidants as described above, were washed twice in M9, and then stained for 2 hours in 1 mL of M9 containing 150 μM 2’,7’-dichlorofluorescein diacetate (DCFDA, Sigma) while rotating in the dark. Worms were then washed twice with M9, and transferred to 2% agarose pads on glass slides, covered, and immediately imaged within 30 minutes of washing out the DCFDA.^48,49^ Imaging was performed as previously described. Images were analyzed in ImageJ, where average intensity of the head region was scored.

### Survival assay and lifespan curves

To measure survival of animals to acute oxidative stress, 50 worms, exposed to oxidants as described above, were washed twice in M9, and then transferred to a 24-well plate containing 1 ml M9 with 250 μM juglone. Animals observed as rigid and immobile after imparting movement to the liquid in the well were scored as dead. Animals were scored for survival every 30 minutes for the first 2 hours and every 1 hour thereafter until all animals were scored as dead.^4,45^ Lifespan curves and statistical analysis of mean lifespan were performed using the Online Application for Survival Analysis 2 (OASIS 2).^50^

### Statistical analysis

Data was compiled and analyzed using JMP Pro v.14.2. The standard least squares second order Scheffe model accounting for the individual components, mixture interactions, and effect of the process variables was used.^51^ The block effect in the first design and the split plot in the second design were taken to be random effects. All effects, including interactions between the mixtures and process variables, with p value smaller than 0.05 during statistical testing (α < 0.05) were considered as significant. Since this approach means dealing with multiple *C. elegans* populations (a population is considered animals cultured on the same NGM plate), we monitored both the mean and variability of gene expression to ensure that population responses are within statistical limits of each other. To assess whether different populations were comparable, we used control limits. A population plotted within the control limits is equivalent to failing to reject the null hypothesis of statistical control (i.e., all populations have equal means), and a population plotted outside the control limits is equivalent to rejecting the null hypothesis. The process mean was monitored using the X bar chart, while process variability was monitored using the S chart.^52^ The responses of the controls were plotted using an Xbar (average) and S (standard deviation) control chart (**Figure S1**).

## Results

### Dose dependency of *gst-4* response

To identify relevant concentrations of chemical mixtures of plumbagin, 1, 4-naphthoquinone, and juglone, (**Figure 1A, B, C**) we first determined the dose-dependent *gst-4* activity to individual components (**Figure 1D**). Animals were exposed for 8 hours, based on prior studies that suggest this time is sufficient to observe a *gst-4* response.^53,54^ *gst-4* belongs to a class of enzymes used to catalyze conjugation of glutathione with xenobiotics. Prior work determined that *gst-4* expression is increased in animals stressed with xenobiotics, while external ROS generated by hypoxanthine/XOD system, UV light, and heat did not elicit a response.^40^ As expected, we identified that as the naphthoquinones concentration increases, *gst-4* activity also increases and eventually saturates (**Figure 1D**), as described in previous studies.^53,54^ To assess the effect of mixture proportions on oxidative stress response, the total naphthoquinone dosage should remain constant. Based on these results, we fixed a combined total dosage of 30 μM for mixture experiments, which allows studying the interactions of components at different proportions without potential saturation of the *gst-4* response. Sublethal doses in the range of 20-30 μM for the 3 compounds under study are known to drive SKN-1-dependent expression of *gst-4*, and also induce a hormetic effect driven by SKN-1.^4,18,37^

### Naphthoquinone mixtures show antagonistic effects under *ad libitum* feeding and physiological temperature conditions

The first mixture experiments were performed under *ad libitum* feeding and no heat stress conditions based on the simplex centroid design. The average and standard deviations of the baseline *gst-4* responses of controls run for each of the ten sets of experiments were compared to determine if runs were comparable to each other. All measurements are within statistical control (**Figure S1A**). We used a Scheffe model fit to develop a response surface (**Figure 2A**). The block effect added in the model indicates that populations 4, 6, and 10 exhibited a significant effect (**Table S1**), suggesting some experimental variation for these populations. However, these deviations caused by the block effects are accounted for in the model. The response surface indicates that the individual naphthoquinones induce a higher *gst-4* response than binary and ternary mixtures, suggesting an antagonistic interaction (**Figure 2A, Table S1**). Since animals were maintained at 25 °C and shifted to 20 °C for chemical exposure, a control exposure experiment at 25 °C was tested to determine if the temperature shift could play a role in the observed responses. The control experiment maintained at 25 °C shows the same *gst-4* expression trend as those exhibited by animals shifted to 20 °C for exposure (**Figure S2 and Figure 2A**): naphthoquinone mixtures are antagonistic in driving the *gst-4* response.

**Figure 2:**
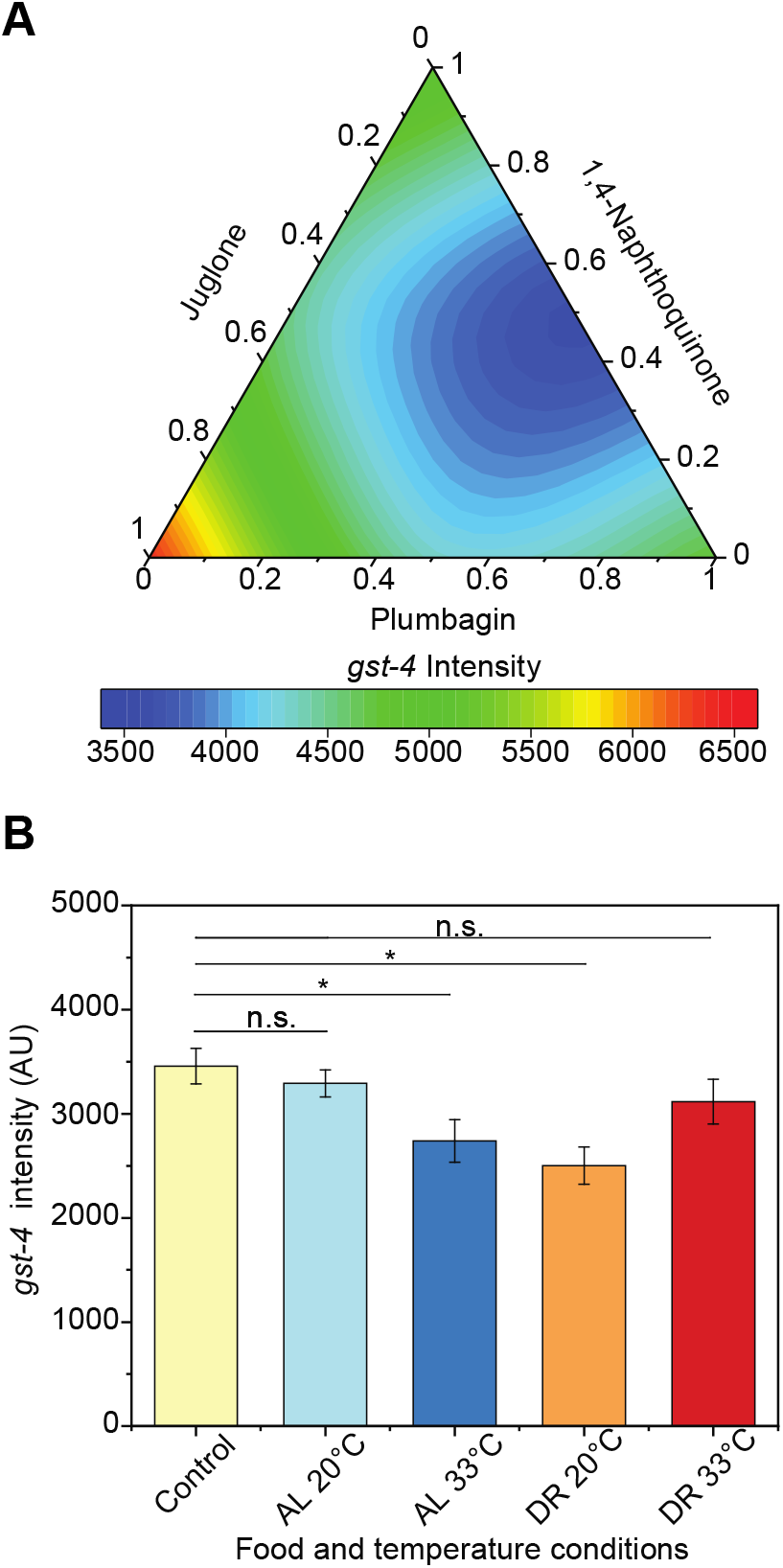
Naphthoquinone mixtures show no synergistic effects under *ad libitum* feeding and no heat stress conditions. A) Response surface of *gst-4* expression levels in CL2166 animals under oxidative stress, 20°C, and *ad libitum* feeding. Response surface modeled using standard least squares second order Scheffe model where main effects and interactions were tested for significance (Table S1). B) Testing for main effect of temperature and food concentration. p > 0.05 (n.s.), p < 0.05 (*). p-values were calculated using Dunnett’s test with *ad libitum*, overall control as control after two-way ANOVA comparison in JMP 14.2. Error bars are SEM. A control at 25 °C is represented as a bar plot in Figure S2.

### Naphthoquinone mixtures induce different *gst-4* responses under different environmental conditions

Temperature and dietary intake have been shown to affect the oxidative stress response in *C. elegans*.^33,37,55^ Thus, we tested how exposure to heat stress and dietary restriction modified the *gst-4* response to naphthoquinone mixtures by fitting the *gst-4* expression data to a mixed effects model. The overall control was plotted as Xbar and S chart and indicates that the populations are within statistical limits of each other (**Figure S1B**). Block effects analysis revealed population 5 exhibited a statistically significant effect compared to the other 27 populations (**Table S2**). However, the model corrects for the effect caused by the treatment application process on population 5.

Heat stress and dietary restriction (DR) significantly affect the oxidative stress response, inducing a lower baseline *gst-4* response than a control of 20 °C and *ad libitum* conditions (**Figure 2B**). However, heat and dietary restriction induce differential responses in *gst-4* in the presence of naphthoquinone mixtures (**Figure 3A, B**). The *ad libitum* and 20 °C response surface (**Figure 3B**) is the repetition of the first mixture experiment and was confirmed to not be statistically different (**Figure 2A**). The experimentally acquired data is represented as conventional bar plots in **Figure S3**. As the temperature is increased to 33 °C, heat stress inhibits the *gst-4* response to pure components and mixtures, consistent with prior results.^37^ Dietary restriction reduces the baseline *gst-4* level (**Figure 2B**).^56^ In contrast, reduced food concentration did not affect the *gst-4* response to 1,4-naphthoquinone and plumbagin (**Figure 3B**) and it drastically increased the response to juglone, as compared to *ad libitum* conditions. Furthermore, exposure to binary or ternary mixtures under dietary restriction did not result in antagonistic interactions observed in mixtures at *ad libitum* dietary regime (**Figure 3B, C**). These results suggest that dietary restriction can differentially modulate the oxidative stress response to individual compounds, and significantly modify interactions amongst naphthoquinones.

**Figure 3:**
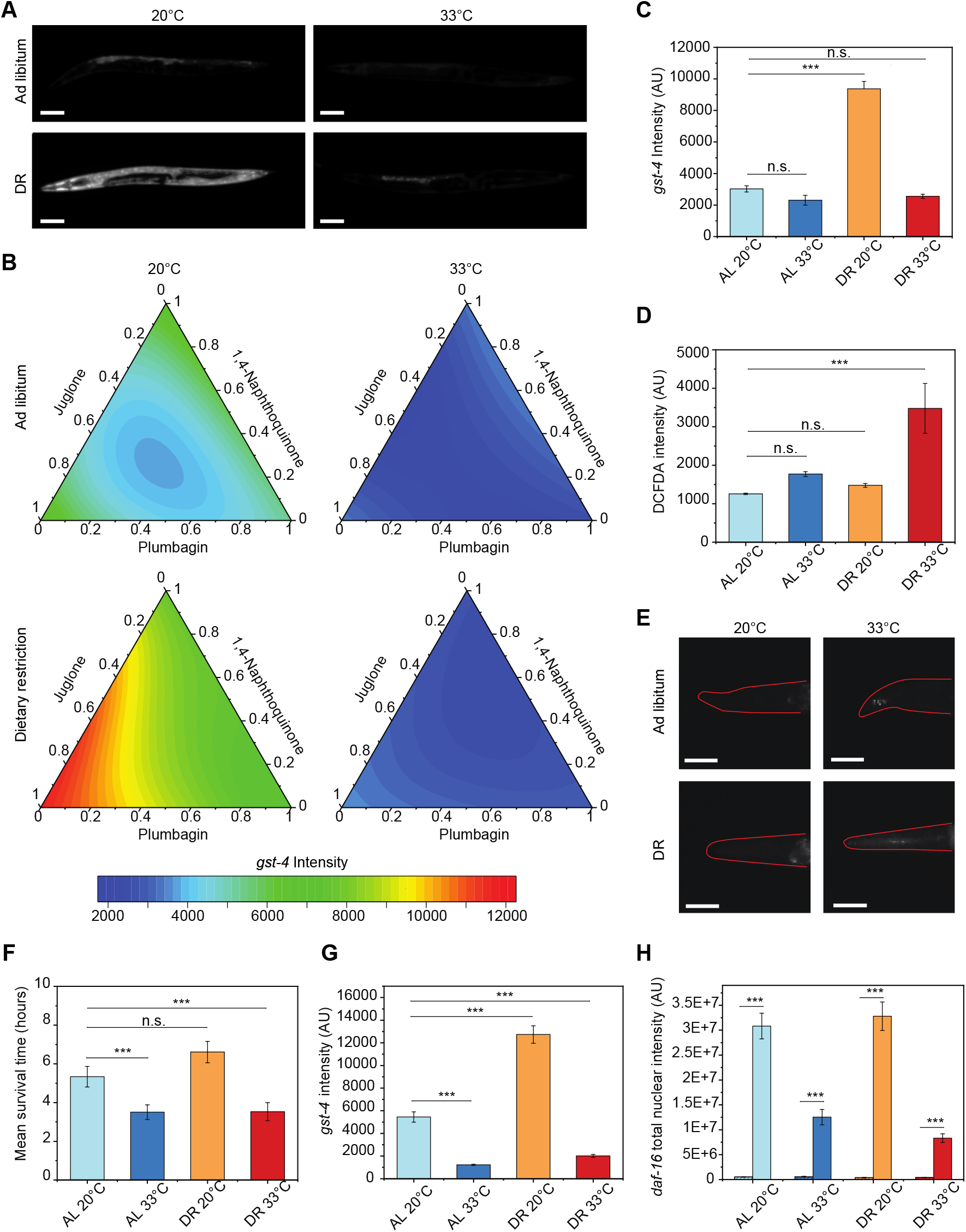
Naphthoquinone mixtures elicit differential response under simultaneous stress exposure. A) Representative CL2166 animals exposed to ternary naphthoquinone mixtures (1,4-Naphthoquinone, Juglone, Plumbagin) at different environmental conditions. B) Response surface of *gst-4* expression levels for CL2166 animals under oxidative stress at different environmental conditions. C) *gst-4* expression levels for CL2166 animals under exposure to the ternary naphthoquinone mixture. D) DCFDA intensity levels for N2 animals exposed to the ternary naphthoquinone mixture. E) DCFDA staining of representative N2 animals exposed to the ternary naphthoquinone mixture. F) Average survival time under exposure to 250 μM Juglone in CL2166 animals pre-exposed to the ternary naphthoquinone mixture at different environmental conditions. G) *gst-4* expression levels under *daf-16* RNAi for CL2166 animals exposed to ternary naphthoquinone mixtures. H) *daf-16* expression levels for MAH97 animals exposed to ternary naphthoquinone mixtures, measured as total intensity in the nuclei of cells per worm. AL: *ad libitum.* DR: dietary restriction. p > 0.05 (n.s.), p < 0.05 (*), p < 0.001 (***). p-values were calculated using Dunnett’s test with *ad libitum*, 20 °C as control after one-way ANOVA comparison in JMP 14.2.Response surfaces modeled using standard least squares second order Scheffe model where main effects and interactions were tested for significance (Table S2). Scale bars are 100 μm. Error bars are SEM. The experimentally acquired data is represented as conventional bar plots in Figure S5.

We next asked whether these environmental modifications of the oxidative stress response to naphthoquinone mixtures could stem from differences in ROS levels, and if the *gst-4* response exhibiting differential responses to mixtures and environmental variables would imply differences in organismal resistance to acute oxidative stress. These questions were addressed by measuring ROS levels through DCFDA staining of N2 animals (**Figure 3D, E**) and by assessing the survival of worms to acute oxidant exposures after exposure to low-level naphthoquinone mixtures (**Figure 3F, S4**). These experiments show that the presence of ROS in the worms is highest under conditions of dietary restriction at 33 °C, which are also conditions with the lowest level of *gst-4* (**Figure 3C**). The differences in ROS levels between the populations exposed to heat stress could be explained by an increase formation of ROS in animals under glucose restriction,^57^ while high levels of glucose renders *C. elegans* more resilient to oxidative stress.^58^ In this case, a parallel can be drawn between glucose and food availability. Animals exposed to acute juglone concentrations exhibit the lowest survival under heat stress (**Figure 3F**), which matches with the lowest observed levels of *gst-4* (**Figure 3C**). These results suggest that environmental conditions that modulate the *gst-4* responses to naphthoquinone mixtures similarly affect the animal’s resistance to oxidants, and that ROS levels are also increased in conditions that result in minimal *gst-4* expression and highest susceptibility to acute juglone exposures.

Since dietary restriction modifies the *gst-4* responses to naphthoquinones (individually and in mixtures), we then asked whether this effect could be modulated by the insulin/insulin-like signaling (IIS) pathway that can be activated with specific dietary restriction regimes.^59^ To address this question, we tested a possible dependence of the *gst-4* response on *daf-16,* the main regulator of the IIS pathway (**Figure 3G, S5**).^60,61^ Comparing the *gst-4* responses in the presence (**Figure 3C**) and absence (**Figure 3G**) of DAF-16 shows that the DR-dependent induction of *gst-4* under naphthoquinone exposures is independent of DAF-16, suggesting an alternative pathway is at play. Likely *skn-1* itself regulates this interaction, since it is known to play a role in DR-induced lifespan extension.^62^ We also measured *daf-16* responses to naphthoquinones mixture exposure by assessing the levels of a DAF-16::GFP fusion protein within intestinal cell nuclei. Like *gst-4, daf-16* responses are inhibited by heat stress (**Figure 3H**). On the other hand, ternary naphthoquinone mixtures induced strong *daf-16* responses in both DR and AL conditions. As mentioned above, under AL conditions, *gst-4* exhibits a reduced signal for ternary mixtures (i.e., antagonistic interactions), which is not observed in *daf-16* activity. This result suggests naphthoquinone mixtures induce strong *daf-16* activity, which is inhibited by heat stress, and the *daf-16* response to oxidants is not modulated by DR, potentially indicating that oxidative stress is prioritized over food intake for downstream signaling.

## Discussion

In this work, we took advantage of a DoE approach to study the oxidative stress response driven by combinatorial exposures to naphthoquinones under a variety of environmental conditions. Unlike traditional full-factorial designs, which would entail an unfeasible number of experiments, using block effects and a simplex centroid design enabled performing experiments in a non-simultaneous and feasible manner. Initially, using mixed second order Scheffe model statistical analysis, we built response curve to ternary naphthoquinone mixtures under controlled environmental factors. The observed antagonistic interactions in **Figure 2A** could stem from different mechanisms of toxicity elicited by the mixture components. For instance, there are differences in the chemical reactivity of juglone and plumbagin.^63^ Naphthoquinones have been shown to result in toxicity through ROS generation or glutathione depletion.^16,18^ Juglone has relatively higher chemical activity and could undergo Michael’s addition to glutathione even at lower doses.^63,64^ 30 μM of 1, 4-naphthoquinone and plumbagin could cause oxidative stress through redox cycling at lower doses, and only at higher doses cause Michael’s addition to glutathione.^63,65^ Mixtures of these chemicals could thus result in lower *gst-4* activation than individual mixtures, if *gst-4* induction is more sensitive towards one of these toxicity mechanisms.

We further analyze if the response to naphthoquinones would be modified under different environmental conditions: food availability and temperature, and found drastic changes to response curves. The reduction in *gst-4* expression levels by heat stress observed in **Figure 2B** could be explained by organismal prioritization of the heat shock response over the oxidative stress response, as previously suggested by Crombie et al.^37^ The reduction in *gst-4* levels by dietary restriction could be the result of the reduced activity of *cct-4* under dietary restriction, which encodes a chaperonin directly involved in SKN-1–dependent transcription of *gst-4*.^56^ We also built response curves to ternary naphthoquinone mixtures under the four environmental conditions, which revealed significant interactions between chemicals. Surprisingly, these interactions were drastically modulated by environmental conditions of temperature and food availability. The identified *gst-4* responses to naphthoquinones is abolished by heat stress, which could be attributed to prioritization of proteostasis over detoxification, where the HSF-1 driven heat stress response genes are upregulated to prevent protein misfolding.^37^

Dietary restriction also modulated stress response. Under *ad libitum* condition, individual components drove *gst-4* expression to similar levels, however, dietary restriction-induced oxidative stress response show high specificity for juglone, suggesting differences in organismal processing of similar oxidants. Although there is cross-regulation between the diet regulated DAF-16 and oxidative stress regulated SKN-1 pathways,^66,67^ RNAi experiments revealed that *gst-4* responses to mixtures are *daf-16-*independent. However, SKN-1 is also known to modulate DR-induced modulation of longevity,^62^ and is thus likely integrating signals for DR and oxidative stress and driving the observed interactions. Interestingly, *daf-16* is activated by naphthoquinone mixtures in the absence of heat stress, recapitulating the *gst-4* inhibition by heat stress. In contrast, DR did not elicit higher *daf-16* activation than AL conditions, suggesting organismal responses to oxidants are prioritized over reduced caloric intake. It is still unclear why the DR effects are specific to the oxidant type. Potentially, these differences could stem from differences in toxicity and detection mechanisms through chemosensation, as explained before, coupled with multiple transcriptional pathways interacting at the organismal level.

These findings highlight the importance of experimental analysis in realistic settings, where a variety of chemical components is present, and where environmental conditions vary significantly. In addition, the identified interactions between naphthoquinone mixtures, heat stress, and dietary restriction, shed light on organismal integration and processing of stressors and environmental factors. This could stem from differences in oxidant detection in *C. elegans.* For instance, low levels of H_2_O_2_ activate the I2 neuron, while paraquat only elicits a response at a very high concentration.^68^ The difference in the *gst-4* response to individual compounds and mixtures could stem from the combined effects of differences in xenobiotic detection and the mechanisms of toxicity by the different naphthoquinones. This result indicates that individual components do not act through a singular mechanism, and that mixtures can drive significantly different organismal responses than individual components, even for highly similar chemical species. A detailed investigation on neuronal SKN-1 could help elucidate whether the interaction mechanisms between chemical mixtures and environments involve neuronal detection.

*C. elegans* are complex in nature and many factors could affect the response to oxidative stress, such as variation in developmental period, food availability on plate, temperature, as well as biological stochasticity. It is necessary to control and account for such effects as combinatorial mixture experiments cannot be performed using a single population. Control charts have proven to be useful in identifying potential problematic populations, which might show a different response. Such populations can have different biological activity and can affect the results of the experiments being performed. Adding block factors and split plots as random effects to the model also help us compare between populations, even if these exhibit differences that could come from experimental or biological variation. Thus, using a combination of mixture experiments and control charts we have highlighted the differential *gst-4* response by *C. elegans* to oxidative stressor mixtures under different environmental conditions. These results warrant further investigation of the different transcriptional pathways involved and shed light on how organisms respond to variable environments and realistic chemical exposures.

---

**Figure S1:**
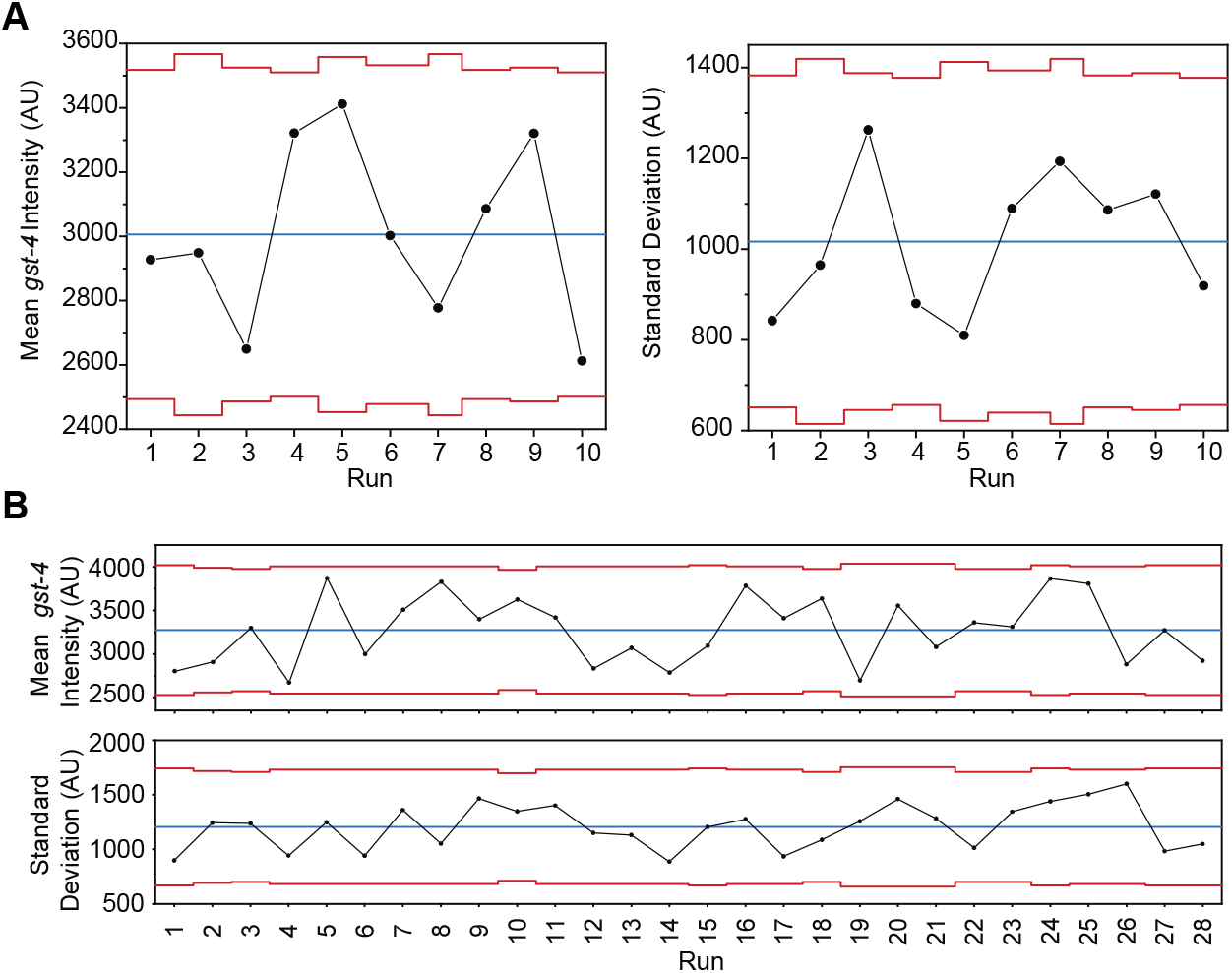
Xbar and S-charts for populations tested. A) Xbar and S-chart for 10 populations used for oxidative stress assays. B) Xbar and S-chart for 28 populations used for oxidative stress assays.

**Figure S2:**
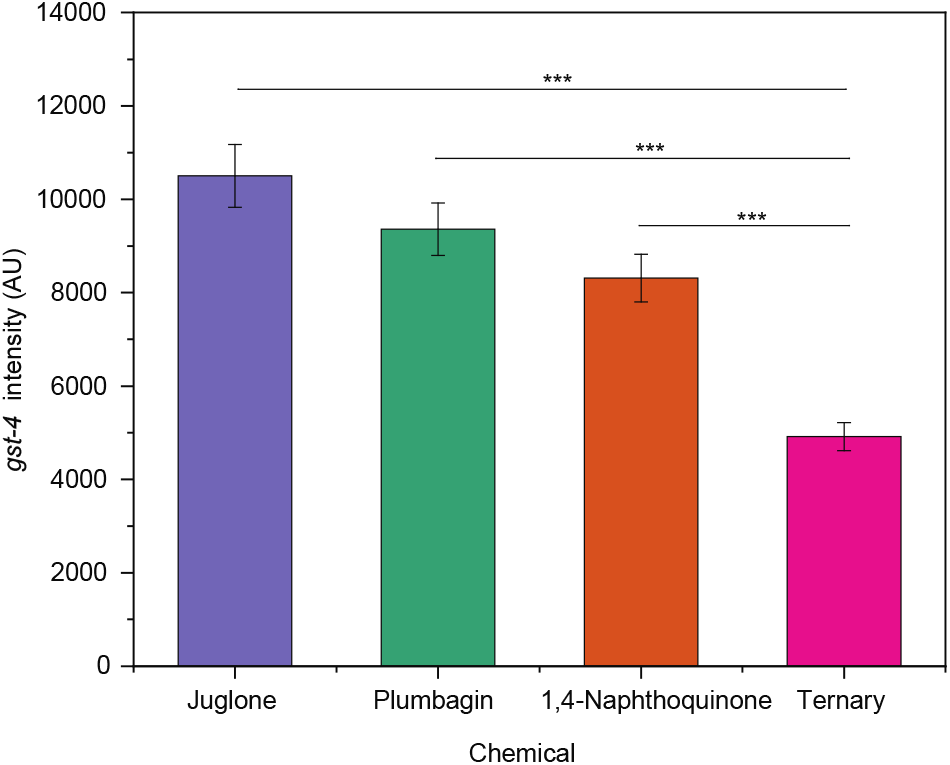
*gst-4* response at 25 °C. *gst-4* response to Plumbagin, 1, 4-Naphthoquinone, Juglone, and ternary mixture at 25 °C. p < 0.001 (***). Values follow the same trend as Figure 2B with a lower value for the ternary mixture (middle point of response surface in Figure 2B), and higher values for individual naphthoquinones (vertex points of response surface in 2B). Error bars are SEM.

**Figure S3:**
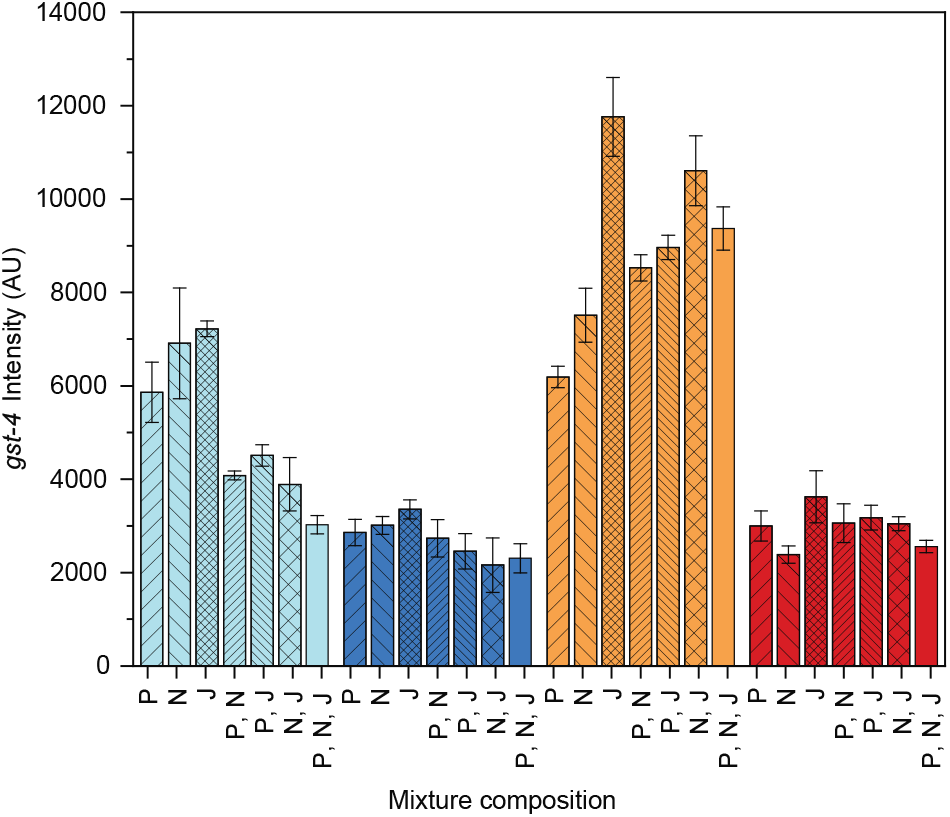
Bar plot representation of experimentally acquired *gst-4* expression level. *gst-4* expression levels of animals exposed to naphthoquinone mixtures at different process conditions. From left to right, AL 20°C (light blue), AL 33°C (blue), DR 20°C (orange), DR 33°C (red). P-Plumbagin, N-1,4-Naphthoquinone, J-Juglone. Error bars are SEM.

**Figure S4:**
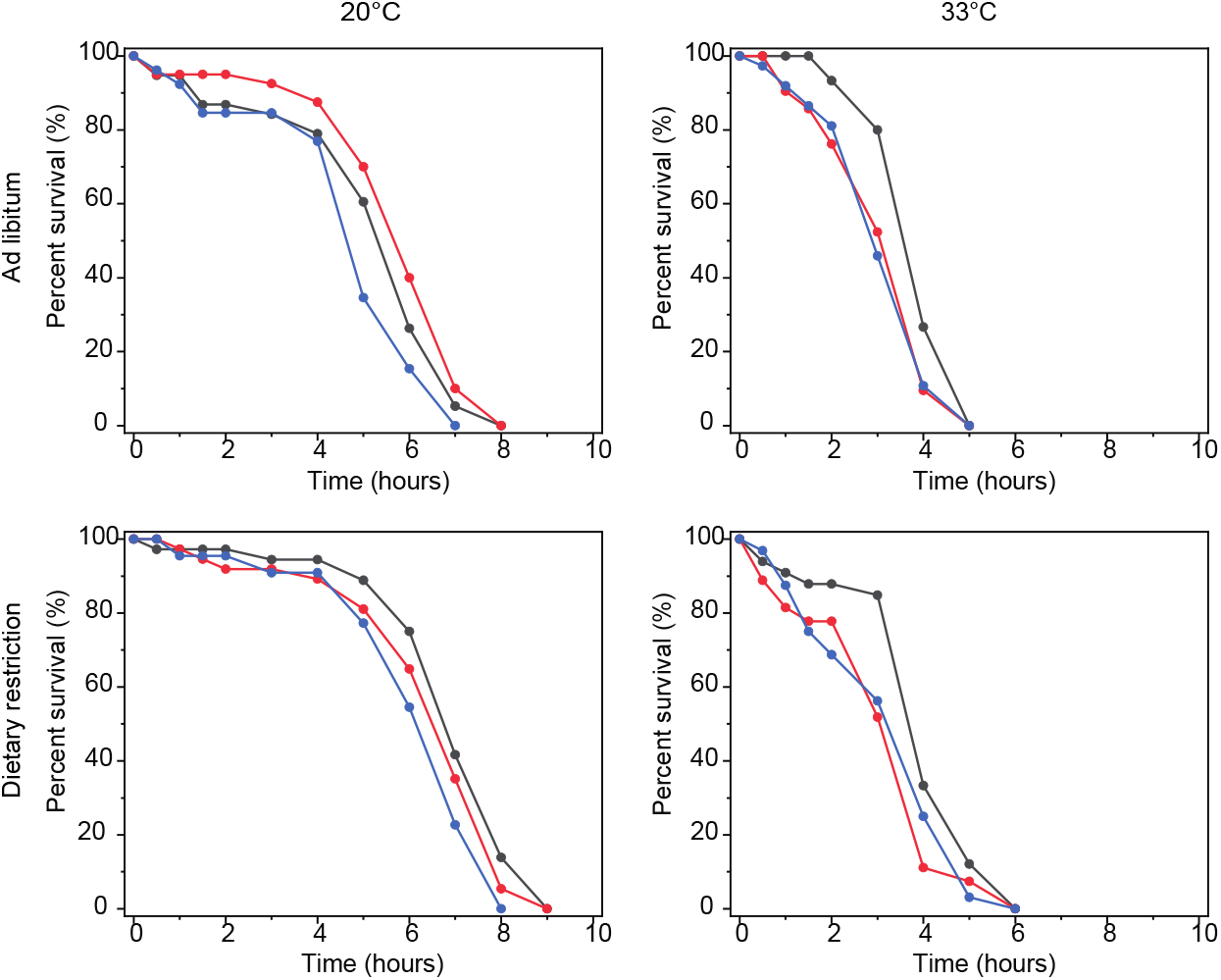
Lifespan curves for juglone survival assay. Lifespan curves built with OASIS 2 for CL2166 animals under ternary mixture at different environmental conditions. Each condition was tested three times.

**Figure S5:**
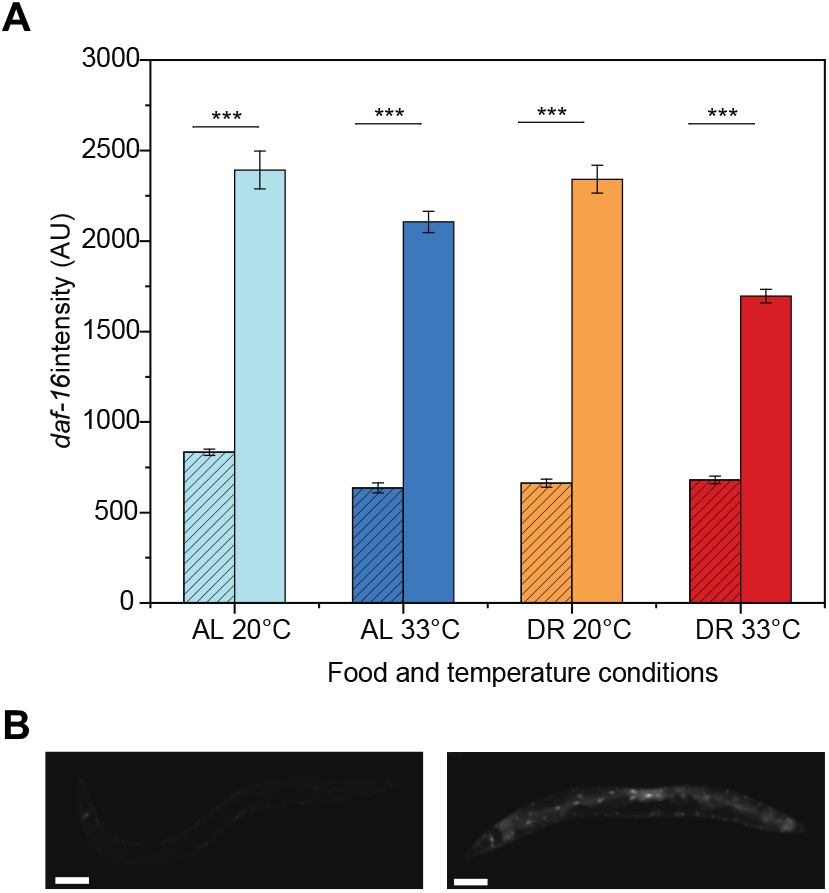
*daf-16* response to ternary naphthoquinone mixture. A) *daf-16* response to ternary mixture at different environmental conditions. Striated bars represent MAH97 animals under *daf-16* RNAi, clear bars are control animals. B) Representative MAH97 animals under daf-16 RNAi (left) and control (right). p < 0.001 (***). Scale bars are 100 μm. Error bars are SEM.

## Acknowledgments

Some strains were provided by the CGC, which is funded by NIH Office of Research Infrastructure Programs (P40 OD010440).

## Data availability

All data underlying this work can be requested from the authors.

## Author contributions

All authors designed the experiments and conceived the work. KSA and JH carried out experiments and analyzed data. JWS and KSA performed design of experiments and statistical analysis. KSA, JH, and ASM wrote the manuscript.

