## Supplementary Tables for "Antagonistic effects of chemical mixtures on the oxidative stress response are silenced by heat stress and reversed under dietary restriction"

**Table S1: Random effect prediction of population blocks and parameter estimates of main effects and interactions for naphthoquinone mixtures under ad libitum conditions.**

| *Term* | *BLUP/Estimate* | *Std Error* | *DFDen* | *t Ratio* | *Prob>\|t\|* |
|---|---|---|---|---|---|
| Population Block[1] | 275.37 | 321.85 | 20.79 | 0.86 | 0.402 |
| Population Block[2] | 300.83 | 322.62 | 20.75 | 0.93 | 0.3618 |
| Population Block[3] | -73.13 | 321.92 | 20.79 | -0.23 | 0.8225 |
| Population Block[4] | 724.31 | 322.62 | 20.75 | 2.25 | 0.0358 |
| Population Block[5] | 332.86 | 322.62 | 20.75 | 1.03 | 0.3141 |
| Population Block[6] | -1076.84 | 322.60 | 20.81 | -3.34 | 0.0031 |
| Population Block[7] | 161.88 | 321.13 | 20.77 | 0.50 | 0.6195 |
| Population Block[8] | 165.11 | 321.85 | 20.79 | 0.51 | 0.6133 |
| Population Block[9] | -119.89 | 321.43 | 20.75 | -0.37 | 0.7129 |
| Population Block[10] | -690.51 | 322.96 | 20.79 | -2.14 | 0.0446 |
| Juglone(Mixture) | 6647.05 | 339.80 | 29.88 | 19.56 | <.0001 |
| 1,4-Naphthoquinone(Mixture) | 5221.82 | 339.80 | 29.88 | 15.37 | <.0001 |
| Plumbagin(Mixture) | 4767.03 | 316.88 | 27.64 | 15.04 | <.0001 |
| Juglone*1,4-Naphthoquinone | -7270.80 | 1308.75 | 25.49 | -5.56 | <.0001 |
| Juglone*Plumbagin | -4855.49 | 1262.85 | 25.02 | -3.84 | 0.0007 |
| 1,4-Naphthoquinone*Plumbagin | -6065.19 | 1269.63 | 25.14 | -4.78 | <.0001 |
| Juglone*1,4-Naphthoquinone*Plumbagin | 10021.04 | 8789.28 | 25.07 | 1.14 | 0.265 |

**Table S2: Random effect prediction of split plots and parameter estimates of main effects and interactions for naphthoquinone mixtures under simultaneous stress exposure.**

| *Term* | *BLUP/Estimate* | *Std Error* | *DFDen* | *t Ratio* | *Prob>\|t\|* |
|---|---|---|---|---|---|
| Whole Plots[1] | -575.96 | 450.16 | 14.27 | -1.28 | 0.2211 |
| Whole Plots[2] | 379.51 | 450.61 | 14.13 | 0.84 | 0.4137 |
| Whole Plots[3] | 310.30 | 450.16 | 14.27 | 0.69 | 0.5017 |
| Whole Plots[4] | 79.01 | 444.15 | 15.16 | 0.18 | 0.8612 |
| Whole Plots[5] | 986.22 | 445.56 | 15.25 | 2.21 | 0.0425 |
| Whole Plots[6] | 141.25 | 450.27 | 14.22 | 0.31 | 0.7583 |
| Whole Plots[7] | -26.32 | 446.61 | 14.85 | -0.06 | 0.9538 |
| Whole Plots[8] | 210.75 | 446.53 | 14.89 | 0.47 | 0.6438 |
| Whole Plots[9] | 79.75 | 444.79 | 15.02 | 0.18 | 0.8601 |
| Whole Plots[10] | -58.62 | 447.60 | 14.83 | -0.13 | 0.8976 |
| Whole Plots[11] | -290.92 | 450.17 | 14.24 | -0.65 | 0.5284 |
| Whole Plots[12] | -212.05 | 447.65 | 14.82 | -0.47 | 0.6426 |
| Whole Plots[13] | -433.91 | 446.53 | 14.89 | -0.97 | 0.3467 |
| Whole Plots[14] | -164.06 | 450.55 | 14.14 | -0.36 | 0.7212 |
| Whole Plots[15] | -147.08 | 453.01 | 13.66 | -0.32 | 0.7503 |
| Whole Plots[16] | 211.95 | 448.67 | 14.44 | 0.47 | 0.6437 |
| Whole Plots[17] | -57.95 | 476.77 | 12.05 | -0.12 | 0.9053 |
| Whole Plots[18] | 82.79 | 445.56 | 15.25 | 0.19 | 0.855 |
| Whole Plots[19] | -351.15 | 445.56 | 15.25 | -0.79 | 0.4427 |
| Whole Plots[20] | 296.32 | 446.92 | 14.86 | 0.66 | 0.5175 |
| Whole Plots[21] | -536.29 | 449.37 | 14.33 | -1.19 | 0.2521 |
| Whole Plots[22] | -645.85 | 447.61 | 14.83 | -1.44 | 0.1698 |
| Whole Plots[23] | -379.74 | 447.90 | 14.75 | -0.85 | 0.4101 |
| Whole Plots[24] | 360.88 | 448.96 | 14.40 | 0.80 | 0.4346 |
| Whole Plots[25] | 650.35 | 443.89 | 15.14 | 1.47 | 0.1633 |
| Whole Plots[26] | -450.20 | 447.77 | 14.81 | -1.01 | 0.3308 |
| Whole Plots[27] | -91.28 | 446.59 | 14.93 | -0.20 | 0.8408 |
| Whole Plots[28] | 632.30 | 453.01 | 13.66 | 1.40 | 0.185 |

| | | | | | |
|---|---|---|---|---|---|
| Juglone(Mixture) | 6447.75 | 295.31 | 55.91 | 21.83 | <.0001 |
| 1,4-Naphthoquinone(Mixture) | 5137.20 | 295.83 | 55.96 | 17.37 | <.0001 |
| Plumbagin(Mixture) | 4380.48 | 295.38 | 55.92 | 14.83 | <.0001 |
| Juglone*1,4-Naphthoquinone | -3024.96 | 1375.96 | 46.58 | -2.20 | 0.0329 |
| Juglone*Plumbagin | -2995.59 | 1343.95 | 42.43 | -2.23 | 0.0312 |
| Juglone*Dietary level | -1191.47 | 295.31 | 55.91 | -4.03 | 0.0002 |
| Juglone*Heat Stress level | -2927.53 | 295.31 | 55.91 | -9.91 | <.0001 |
| 1,4-Naphthoquinone*Plumbagin | 429.22 | 1350.27 | 42.74 | 0.32 | 0.7521 |
| 1,4-Naphthoquinone*Dietary level | -14.65 | 295.83 | 55.96 | -0.05 | 0.9607 |
| 1,4-Naphthoquinone*Heat Stress level | -2143.92 | 295.83 | 55.96 | -7.25 | <.0001 |
| Plumbagin*Dietary level | -392.97 | 295.38 | 55.92 | -1.33 | 0.1888 |
| Plumbagin*Heat Stress level | -1607.82 | 295.38 | 55.92 | -5.44 | <.0001 |
| Dietary level*Heat Stress level | 3198809.40 | 3503890.00 | 42.48 | 0.91 | 0.3664 |
| Juglone*1,4-Naphthoquinone*Plumbagin | -9811.49 | 9463.33 | 42.48 | -1.04 | 0.3057 |
| Juglone*1,4-Naphthoquinone*Dietary level | -4610.71 | 1375.96 | 46.58 | -3.35 | 0.0016 |
| Juglone*1,4-Naphthoquinone*Heat Stress level | 69.72 | 1375.96 | 46.58 | 0.05 | 0.9598 |
| Juglone*Plumbagin*Dietary level | -2178.32 | 1343.95 | 42.43 | -1.62 | 0.1125 |
| Juglone*Plumbagin*Heat Stress level | 1264.48 | 1343.95 | 42.43 | 0.94 | 0.3521 |
| Juglone*Dietary level*Heat Stress level | -3197723.00 | 3503871.00 | 42.48 | -0.91 | 0.3666 |
| 1,4-Naphthoquinone*Plumbagin*Dietary level | -2605.30 | 1350.27 | 42.74 | -1.93 | 0.0603 |
| 1,4-Naphthoquinone*Plumbagin*Heat Stress level | -507.70 | 1350.27 | 42.74 | -0.38 | 0.7088 |
| 1,4-Naphthoquinone*Dietary level*Heat Stress level | -3198449.00 | 3503874.00 | 42.48 | -0.91 | 0.3665 |
| Plumbagin*Dietary level*Heat Stress level | -3198617.00 | 3503869.00 | 42.48 | -0.91 | 0.3665 |
| Juglone*1,4-Naphthoquinone*Plumbagin*Dietary level | 3274.24 | 9463.33 | 42.48 | 0.35 | 0.7311 |
| Juglone*1,4-Naphthoquinone*Plumbagin*Heat Stress level | 7196.80 | 9463.33 | 42.48 | 0.76 | 0.4512 |
| Juglone*1,4-Naphthoquinone*Dietary level*Heat Stress level | 2508.55 | 1375.96 | 46.58 | 1.82 | 0.0747 |
| Juglone*Plumbagin*Dietary level*Heat Stress level | 1301.67 | 1343.95 | 42.43 | 0.97 | 0.3383 |
| 1,4-Naphthoquinone*Plumbagin*Dietary level*Heat Stress level | 1424.62 | 1350.27 | 42.74 | 1.06 | 0.2973 |
